# ProGeo-neo: a Customized Proteogenomic Workflow for Neoantigen Prediction and Selection

**DOI:** 10.1101/719351

**Authors:** Yuyu Li, Guangzhi Wang, Xiaoxiu Tan, Jian Ouyang, Menghuan Zhang, Xiaofeng Song, Qi Liu, Qibin Leng, Lanming Chen, Lu Xie

## Abstract

Neoantigens can be differentially recognized by T cell receptor (TCR) as these sequences are derived from mutant proteins and are unique to the tumor. The discovery of neoantigens is the first key step for tumor-specific antigen (TSA) based immunotherapy. Based on high-throughput tumor genomic analysis, each missense mutation can potentially give rise to multiple neopeptides, resulting in a vast total number, but only a small percentage of these peptides may achieve immune-dominant status with a given major histocompatibility complex (MHC) class I allele. Specific identification of immunogenic candidate neoantigens is consequently a major challenge. Currently almost all neoantigen prediction tools are based on genomics data. Here we report the construction of proteogenomics prediction of neoantigen (ProGeo-neo) pipeline, which incorporates the following modules: mining tumor specific antigens from next-generation sequencing genomic and mRNA expression data, predicting the binding mutant peptides to class I MHC molecules by latest netMHCpan (v.4.0), verifying MHC-peptides by MaxQuant with mass spectrometry proteomics data searched against customized protein database, and checking potential immunogenicity of T-cell-recognization by additional screening methods. ProGeo-neo pipeline achieves proteogenomics strategy and the neopeptides identified were of much higher quality as compared to those identified using genomic data only. The pipeline was constructed based on the genomics and proteomics data of Jurkat leukemia cell line but is generally applicable to other solid cancer research. With massively parallel sequencing and proteomics profiling increasing, this proteogenomics workflow should be useful for neoantigen oriented research and immunotherapy.

## 1. Introduction

Cancer neoantigens arise from tumor-specific mutations, which are bound to human leukocyte antigen (HLA) molecules and shuttled to the cell surface, they are highly immunogenic because they are not present in normal tissues and hence bypass central thymic tolerance [1]. In human being, the MHC molecules are encoded by a cluster of genes on chromosome 6 and often referred to as HLA. They are broadly split into two types: MHC-class I (HLA-I) and MHC-class II molecules (HLA-II). MHC class I-associated peptides are generated following degradation of intracellular proteins by the ubiquitin-proteasome system, which can be recognized by cytotoxic CD8+T cells [2]. Helper CD4+T cells recognize MHC class II-associated peptides, which are derived from protease-mediated degradation of exogenous proteins in extracellular origin [3]. We chose to focus our study here on class I CD8+T cell epitopes because CD8+ T cells are the main mediators of naturally occurring and therapeutically induced immune responses to cancer.

Neoantigens could be immunogenic and thus become ideal targets for cancer immunity. Some projects have confirmed that tumor-specific antigens can be targets for checkpoint blockade therapy and personalized vaccine therapy [4, 5]. Anagnostou et al found that neoantigens were relevant targets of initial response to checkpoint blockade in non-small cell lung cancer [6]. T cell responses against tumor neoantigens have been observed after immune checkpoint blockage with ipilimumab [7]. Lu et al demonstrated that adoptive T cell therapy targeting a tumor-specific antigen can mediate long-term survival for a patient with metastatic melanoma [8]. It has also been shown that naive T cells from non-cancer patients could react against tumor neoantigens, providing evidence that de novo anti-neoantigen responses could be elicited [9]. Furthermore, two studies examining the effects of neoantigen vaccines on patients with stage III or IV melanoma demonstrated clinical safety and immunogenicity efficacy data in phase I studies [1, 10].

In recent years, technologies in genomics and proteomics have been significantly improved, in the meantime some supportive bioinformatics and in silico HLA-binding prediction tools have been developed. Multiple immunoinformatics studies endeavored to predict mutation-derived neoantigens and identify those with clinical relevance from large-scale cancer sequencing data [11, 12]. These methods, such as Tumor Immunology miner (TIminer) [13], pVACSeq [14], INTEGRATE-neo [15], TSNAD [16], have contributed to a major breakthrough in the discovery of neoantigens. TIminer integrates bioinformatics tools to predict tumor neoantigens through analyzing single-sample RNA-seq data and somatic DNA mutations. pVACSeq combines the tumor mutation and expression data to predict and filter neoantigens. INTEGRATE-neo was designed to predict neoantigens from fusion genes, which combines peptide prediction and HLA allele prediction results. TSNAD is an integrated software for cancer somatic mutation and tumor-specific neoantigen detection.

However, previous neoantigen prediction tools only predict HLA-binding with genomic and transcriptome data, without considering proteomics data. In this work, we constructed a workflow to predict and verify HLA-I binding peptides on personalized level by proteogenomic strategy which integrates genomics and proteomics data. The workflow was developed based on a specifed tumor cell line data, but it would be applicable to specific tumor patient data. The workflow is implemented in a software package, ProGeo-neo. ProGeo-neo integrated latest genomics and proteomics data analysis as well as neoantigen prediction methods, with in-house coding, and further filtering and immunogenicity screening, to facilitate users to predict and select tumor neoantigen. ProGeo-neo consists of three modules: construction of customized protein sequence database, HLA alleles prediction, neoantigen prediction and filtration. Users can run ProGeo-neo by performing command lines under the Linux operation system (centos6). Full source code and installation instructions are freely available from https://github.com/kbvstmd/ProGeo-neo.

## 2. Material and Methods

The ProGeo-neo workflow construction is illustrated in Figure 1, including: Transcriptome and genome data processing and annotating of mutant peptides; NetMHCpan[17] predicting neoantigens based on peptide-MHC binding affinity; mutant peptides filtered considering transcripts per million (TPM) to select only candidate neoantigens arising from expressed genes; proteogenomics identification of potential predicted neoantigens based on constructed mutant peptidome database and MaxQuant[18] software. Furthermore, sequence similarities between the neoantigens and the cross-reactive microbial peptides were calculated to select more likely immunogenic neoantigens. At last, the workflow was implemented in a software package for potential further application on personalized proteogenomic discovery of tumor neoantigens.

**Figure 1.**
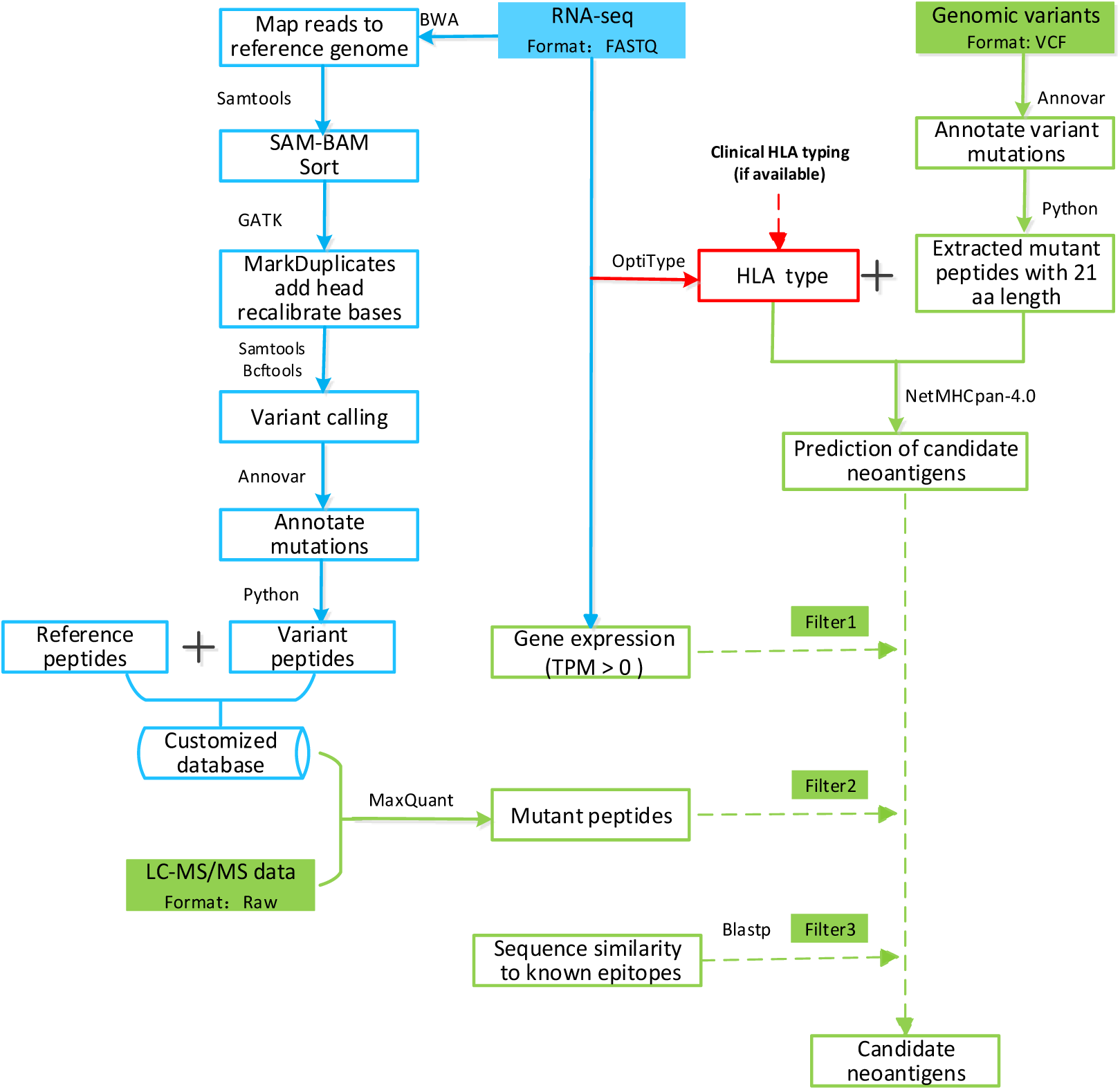
The workflow of ProGeo-neo: neoantigen prediction and selection with proteogenomic strategy. It includes three functional blocks: (1) RNA-seq data analysis, leading to variant peptides, which was used to create a customized database (highlighted in blue above); (2) HLA alleles were inferred from RNA-seq data (highlighted in red above). To increase the flexibility of ProGeo-neo, users have the option to upload their own HLA alleles; (3) Neoantigen prediction based on genomic data. Neoantigen screening by proteomics data (LC-MS/MS). Neoantigen filtration by RNA expression and by T cell receptor recognition (epitope). (highlighted in green above)

### 2.1 Data

The paired end 200 bp sequencing RNA-seq data for the Jurkat cell line generated from Illumina HiSeq 2000 was downloaded at NCBI’s Gene Expression Omnibus (GEO)[19] repository with accession number GSE45428 [20]. LC-MS/MS Jurkat proteomics data generated by LTQ Orbitrap velos in raw format were obtained from PeptideAtlas (http://www.peptideatlas.org/) with the identifier PASS00215. Jurkat whole genome sequencing data were obtained from Zenodo (400615)[21]. The data set of known positive epitopes, that were cross-reactive microbial peptides, was from the Immune Epitope Database[22] (IEDB, http://www.iedb.org/).

Human normal protein sequences were downloaded in fasta format from Uniprot Database(http://www.uniprot.org/)[23].Contaminated protein sequences were downloaded in fasta format from common Repository of Adventitious Proteins (cRAP) (http://www.thegpm.org/crap/).

### 2.2 RNA-seq data processing

The RNA sequencing data in SRA format were converted to fastq with the fastq-dump tool which is part of the SRA Toolkit [24]. Then low-quality reads were removed by using Sickle (version 0.1.18), which confirmed that the sequencing was of high quality. All clean reads were aligned to the human genome (release hg38) using Burrows-Wheeler Alignment tool (BWA) (version 0.7.17) [25]. Samtools (version 1.9) [26] was used to convert the resulting sam files to binary format (bam). Duplicate reads were marked and removed using the Picard tool MarkDuplicates. Base recalibration was performed with GATK (version 4.0.10.1) [27] to reduce false-positive variant calls. Samtools mpileup command was used to calculate the genotype likelihoods supported by the aligned reads. And then the Bcftools (version 1.9) [28] use the genotype likelihoods generated from the previous step to call SNVs, and output all identified variants in the variant call format (VCF). Only single nucleotide variants (SNVs) with a quality score (QUAL) higher than 20 were used for subsequent analysis. Finally, annotation of single nucleotide polymorphisms (SNPs) against hg38 human reference genome was performed using Annovar [29]. Kallisto (V 0.45.0) [30] was used to quantify transcripts expression by TPM level from RNA-seq data.

HLA alleles were inferred from RNA-seq data using OptiType[31] with default settings.

### 2.3 Neoantigen prediction

NetMHCpan has been verified by several groups to be the most accurate tool currently available for predicting neoantigens. Therefore, we integrate it to forecast mutant peptides that bind to MHC class I molecules using artificial neural network algorithm. The predicted HLA alleles and the mutated expressed peptides were used as input for the algorithm NetMHCpan4.0 to estimate their binding affinities and predict neoantigens.

### 2.4 Mutation annotation and peptide extraction

Tumor missense mutations, translated into amino acid substitutions, provide a form of antigens that the immune system perceives as foreign, which elicits tumor-specific T cell immunity. Hence, we focused on missense variants in this study. All the mutations were annotated with Annovar. Sequences corresponding to each of the coding missense mutations that would cause amino-acid substitutions were translated into a 21-mer amino acid fasta sequence, with 10 amino acids flanking the substituted amino acid on each side.

### 2.5 Classification of neoantigen prediction result

Percentile rank scores return a higher sensitivity in MHC ligand identification compared to half-maximum inhibitory Concentration (IC50) [32], the rank score approach starts from the extreme assumption that the number of presented peptides is identical for all MHC molecules, this measure is not affected by inherent bias of certain molecules towards higher or lower mean predicted affinities. We therefore select candidate binders based on %Rank rather than IC50, the smaller of %rank value indicates the stronger affinity of the predicted neoantigen with its corresponding HLA subtype. Each peptide may be classified as a strong binder (%Rank≤0.5), weak binder (0.5<%Rank≤2) or non-binder (%Rank>2). We included both strong and weak binding predicted neoantigens in sebsequent analyses.

### 2.6 Neoantigen filtration

Not all predicted neoantigens resulting from cancer mutations can be expected to express as neo-peptides, or to be immunogenic, as only a small fraction of peptides in most current vaccines are capable of eliciting CD8+T cell responses. It is necessary to screen the reliability of the predicted candidate antigens based on peptide presence and potential immunogenicity.

### 2.7 Identification of mutant peptides in protein level

SNVs from RNA-seq data provide a better reference proteomics datasets compared with WGS, of which the read coverage is generally lower[33]. Therefore, mass spectrometry (MS)-based proteogenomic neoantigen discovery workflow is better to use customized searchable peptide databases derived from tumor RNA-seq. A Python script was written to directly map single amino acid variants (SAAVs) from the Annovar annotated SNPs. Mutant protein sequences with missense mutation sites are generated by substituting the mutant amino acid in human normal protein sequences and all these sequences are appended to the human normal protein and cRAP fasta file. cRAP is a database of protein sequences that are found as contaminants in proteomics experiments.

In this work raw LC-MS/MS Jurkat proteomics spectra were searched against the customized Uniprot+cRAP+Variant peptides database using the MaxQuant to filter neoantigens. MaxQuant identified mutant peptides with MS data, which could verify the presence of expressed neoantigens. The parameter settings of MaxQuant were as follows: for peaklist-generating the default parameters were used; the variable modifications included protein N-terminal acetylation, methionine oxidation; strict trypsin specificity was required allowing up to two missed cleavages; carbamidomethylation of cysteine was set as fixed modification. Reversed sequences were used as a decoy database. False discovery rate (FDR) thresholds for protein, peptides were specified at 1%. Minimum required peptide length was set to 7 amino acids.

### 2.8 Filtering neoantigens potentially recognizable by T cell receptors

An immunogenic peptide should fulfill at least two criteria: presentation by an MHC molecule and recognition by a T-cell receptor. It is known that neoantigen has homology to infectious disease-derived epitopes, which are recognized by the human TCR repertoire. Tumor-infiltrating T cells can cross reactively recognize both cancer neoantigens and homologous non-cancer microbial antigens [34]. The higher the sequence similarity between the neoantigen and the cross-reactive microbial peptide, the greater the probability that the neoantigen is recognized by T cells. Hence, blastp [35] method is used to measure the sequence similarity of neoantigens and cross-reactive microbial peptides in the pipeline. These cross-reactive microbial peptides are linear epitopes from human infectious diseases that are positively recognized by T cells after class I MHC presentation.

## 3. Results

### 3.1 Neoantigens identified by NetMHCpan

Two HLA genotypes of each gene could be obtained because humans are diploid, we only considered mutated epitopes whose HLA class I allele restriction was defined at four-digit resolution, including one HLA-A allele (HLA*A03:01), two HLA-B alleles (HLA-B*07:02, HLA-B*35:03), two HLA-C alleles (HLA*C07:02, HLA*C04:01) identeified from Jurkat cell line.

Class I MHC dimers are responsible for presenting CD8+ cytotoxic T-cell epitopes and binding peptide ligands of 8-11 amino acids. Hence, the results from NetMHCpan were peptides of 8-11 amino acids in length for Jurkat cell line used in this study. The peptides without mutation sites were removed, which resulted in 36,835 expressed candidate neoantigens predicted by NetMHCpan, originated from 9,817 missense mutations based on Jurkat whole genome sequencing data (Supplementary Table S1). Candidate neoantigens included 9,966 high-affinity peptides (%Rank≤0.5) and 26,869 low-affinity peptides (0.5<%Rank≤2). Neoantigens distribution in binding each HLA allele was shown in Figure 2, with the number of neoantigens varied across genotypes and ranged from 6,175 to 8,499. In addition, we observed that some neoantigens were shared between HLA genotypes, which reflects the flexibility of interaction between antigen peptides and MHC molecules, the same type of HLA-I molecules can selectively identify antigenic peptides with the same or similar anchoring residues, this could facilitate generation of novel vaccines for immunoprophylaxis and immunotherapy. These shared neoantigens only represent identical peptides originated from one gene, they have the ability to combine with different HLA typing. MHC-class I genes have significant population variation, with polymorphisms resulting in amino-acid differences particularly concentrated in the region that binds the processed peptides. This results in different binding strengths to the same peptides being conferred by these individual genotypes, which may lead to differences in response intensity. It indicates that MHC molecules with its polymorphism participate in, as well as regulate immune response.

**Figure 2.**
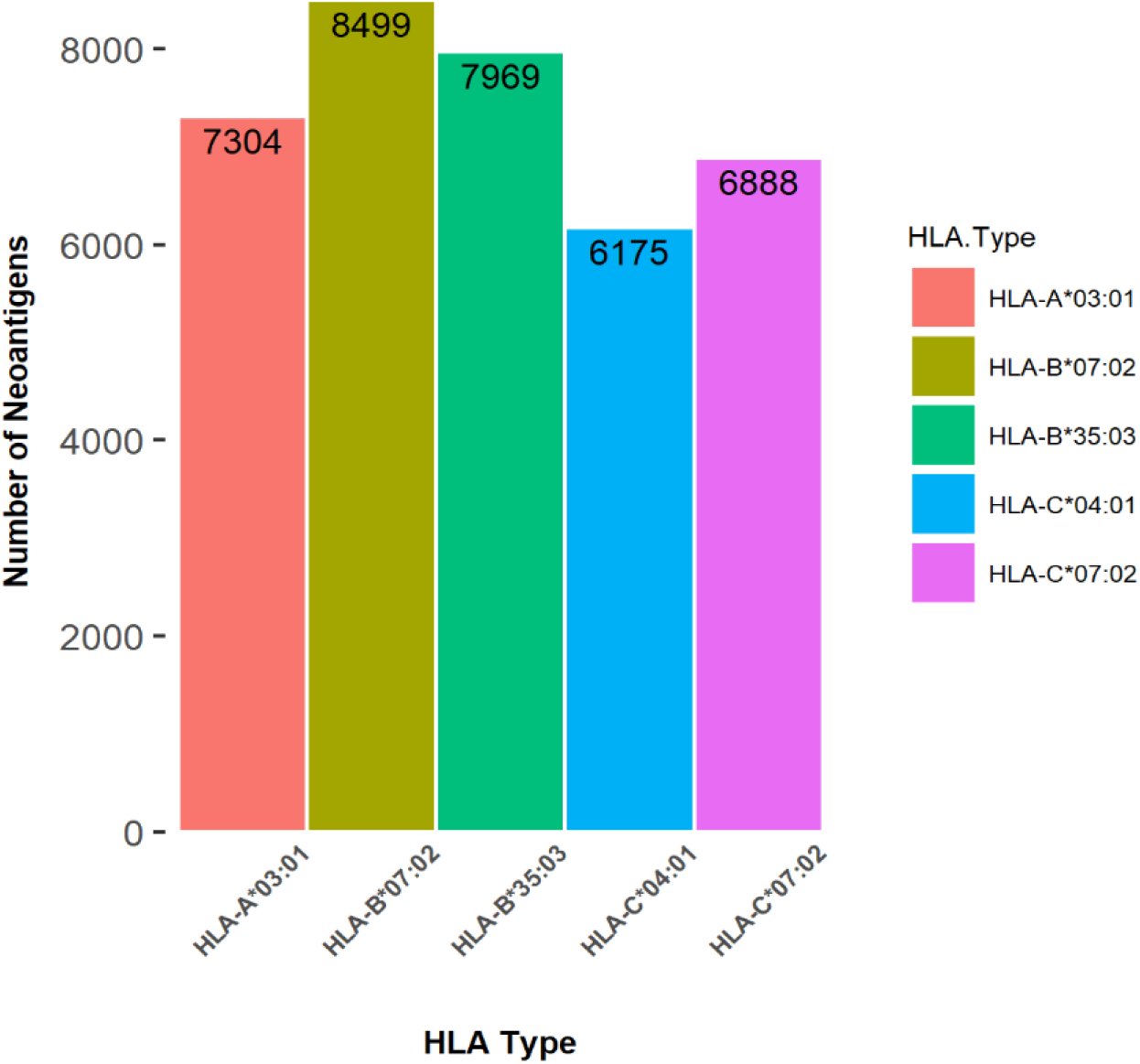
The number of predicted neoantigens binding with each HLA allele

### 3.2 Neoantigens filtered in gene expression level

Peptide presentation is statistically associated with expression of RNA[36, 37]. Peptides from genes expressed at 0 TPM (unexpressed at RNA level) were excluded from candidate peptides (8-11AA) even if their %rank values were≤2, resulted in 30,141 binding peptides being filtered out.

### 3.3 Neoantigens filtered in protein level

MS-based tumor specific antigen discovery workflows must use proteogenomic approaches to build customized databases derived from tumor RNA-seq data [38]. The RNA-seq data were analyzed to find Jurkat cell-specific SNVs and used to create a customized database for peptide identification by MS searching in this study. During the pre-processing of RNA-seq data, only SNVs with a quality score higher than 20 were used for subsequent analysis, which is a phred-scaled score that reflects the confidence of the SNV call. Eventually, high-quality non-synonymous missense mutations were identified. For each missense mutation, a custom Python script was used to translate the reference amino acid to the variant amino acid and totally generated 9,083 mutant protein sequences. Using these mutant protein sequences, along with the human reference proteome (Human 21,410 entries) and cRAP (Human 68 entries), a customized fasta database was created.

The customized database was searched against the MS data using the MaxQuant software with described parameters. 70,622 peptides were identified at a 1% FDR. From these, there were a total of 487 mutant peptides (7-43AA) mapping to 473 missense SNV sites (Supplementary Table S2), corresponding to 0.69% of all peptides. This percentage, representing the proportion of single amino acid polymorphism peptides detected in a shotgun proteomics experiment, is similar to previous findings [39, 40]. We further selected only the candidate neoantigens from the 487 mutant peptides (7-43AA) identified from MaxQuant. Finally, 655 candidate neoantigens with identified peptides proof at protein expression level were retained for further screening of potential clinical relevance (immunogenicity).

### 3.4 Filtering neoantigens potentially recognized by T cell receptors

Peptides of bacterial and viral pathogens can be recognized by immunogenic T cells, therefore neoantigens cross-reactive with these extrogenous pathogens may be recognized by tumor reactive T cells. Among the 655 candidate neoantigens, 313 neoantigens were found to be most likely recognized by the TCR repertoir based on a sequence comparison analysis (Supplementary Table S3), with a comparison probability of 47.79%. The results of sequence similarity between neoantigens and cross-reactive peptides were ranged from 20 to 100. We artificially divided the sequence similarity score of neoantigens into four stages (20-39, 40-59, 60-79, 80-100) (Figure 3), it is observed that the scores of sequence similarity are greater than 60 among the majority of neoantigens. Generally, the higher the sequence similarity, the greater the likelihood that a neoantigen will be recognized by a T cell receptor.

**Figure 3.**
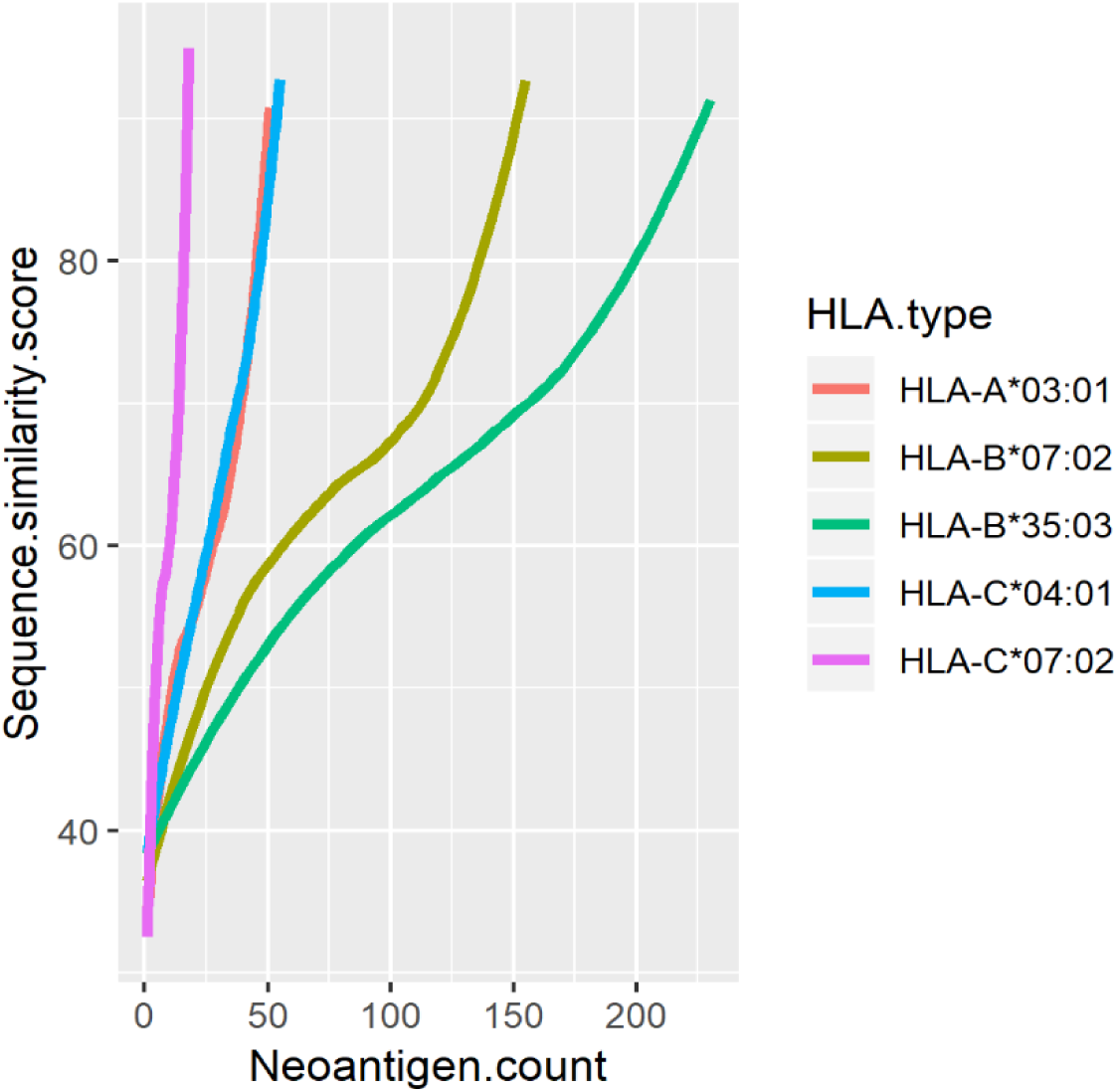
Sequence similarity scores of neoantigens correlating with higher number of cross-reactive microbial peptides.

To test if our method provides the accuracy to identify mutant peptides as neoantigens, we performed the sequence similarity analysis of peptides that were not filtered by mass spectrometry in the same way. It was found that 11,100 peptides have sequence similarity with cross reactive microbial peptides among the 29,486 candidate neoantigens, with a comparison probability of 37.64%, which was significantly lower than our previous alignment probability 47.79%. It proves the validity of identification at the protein level.

### 3.5 Confirmation of potentially significant neoantigens by functional analyses

To acquire understanding of the biological functions of 77 mutant proteins, we used our in-house OmicsBean-Cancer workflow software (http://www.omicsbean-cancer.com/), which incorporated pathway and network analysis tools such as KEGG [41] and protein-protein interactions (PPI) [42]. The results showed that multiple neoantigen proteins participated in AMPK, MAPK, mTOR signaling pathways which are key players in occurrence and development of leukemia from which Jurkat cell line was originated from. Specifically, MAPK14 gene was involved in the regulation of multiple immune signaling pathways, such as Hematopoietic cell lineage, Toll like receptor signaling pathway, etc. MAPK14 can promote the expression of autophagy related genes at transcriptome level, autophagy may lead to the presentation of longer HLA-I peptides from the pathogens or from the self-proteome. Furthermore, the String database was used to depict the integrated PPI of proteins associated with neoantigens, which help to judge the significance of a neoantigen protein by checking its position in PPI network. 55 mutant proteins were interrelated in the neoantigen network (Figure4). We found that eEF2 was a driver gene, and an important node in protein network. It has been reported that eEF2-derived immunogenic peptide was able to further enhance its capacity of inducing antigen-specific cytotoxic T lymphocytes (CTLs) against colon cancer cells [43]. These are examples to show that if the mutant peptides derived from MAPK14 or eEF2 are used as neoantigen targets to stimulate the body, the specific immune response will likely benefit the destruction of tumor cells. These neoantigens may qualify further experimental validation.

### 3.6 Test of other customized data

To test this pipeline, we analyzed the high-throughput sequencing and mass spectrometry data in Mono-allelic Cells [44]. First, we generated a customized personalized reference database based on RNA-seq data. Then, we searched the raw MS data against this database, directly identified 9 peptide ligands harbouring mutations from eluted HLA class I peptides assigned to HLA-B5701. These data of limited size are provided as test data for ProGeo-neo pipeline. Different modules of the pipeline can be called into function according to the availability of data types.

### 3.7 ProGeo-neo: Integrated software for neoantigen prediction and screening based on customized proteogenomics workflow

We have further implemented the proposed approach as an integrated software named ProGeo-neo, which is written in Python programming language (v2.7), and calls for standard third-party software. In order to run normally, third-party softwares are provided, but Licenses for academically used software (NetMHCpan) must be obtained by users. Compared with other neoantigen prediction pipelines, ProGeo-neo has a number of advantages: first, it offers a pipeline for mutation calling from high-throughput sequencing data; second, it offers a proteogenomic strategy to build customized database derived from tumor RNA-seq data; third, it not only takes into consideration the neoantigens presented by class I MHC molecules, but also directly identifies these mutant peptides with MS data; fourth, it offers additional filtration criteria: tumor vs. closest microbial peptide sequence similarity.

The software consists of three toolkits: construction of customized protein sequence database, HLA alleles prediction, neoantigen prediction and filtration. Before running the toolkit, users need to configure the software paths and parameters. This step is of great significance. After setting the configurations, users can run the pipeline by performing command lines. Detailed operations and scripts used to produce this result are provided in the user’s manual. ProGeo-neo is available at https://github.com/kbvstmd/ProGeo-neo.

## 4. Discussion

Based on high-throughput tumor genomic analysis, each missense mutation can potentially give rise to multiple neopeptides, resulting in a vast total number, only one out of 2,000 of the peptides may achieve immunodominant status with a given class I allele[45] (Figure 5A). Specific identification of immunogenic candidate neoantigens is consequently a major challenge. Here we provide ProGeo-neo pipeline, by incorporating state of the art key bioinformatics tools and screening methods in one workflow, the numbers of reliable neoantigen candidates can be greatly cut down (Figure 5B), which will benefit preclinical studies. In our work, 9,871 mutant peptides (21AA), which map to 5,482 genes, give rise to 373,046 theoretical mutant peptides. About 10% potential HLA-I binding neoantigens were identified by NetMHCpan. Tandem mass spectra were processed by MaxQuant to infer mutant peptides, only a minority of them were verified at the level of proteome (Figure 5B). The neopeptides identified from this customized database workflow were of much higher quality as compared to those identified using genomic data without filtering. Finally, sequence homology analysis was performed based on blastp algorithm, 313 neoantigens on 77 mutant proteins were identified. Furthermore, we found that the fraction of neoantigens derived from driver genes were less than the number of neoantigens expressed by passenger genes (Figure 4), which was consistent with previous report [46]. A direct implication of this bias in neoantigen-specific T cell reactivity toward patient-specific passenger mutations is that the targeting of defined neoantigens will likely require the development of personalized immunotherapies [47]. Furthermore, ProGeo-neo can be applied not only to identify putative neoantigens, but also to compare neoantigens with cross-reactive microbial peptides. The analysis results can be used for subsequent clinical biomarker discovery.

**Figure 4.**
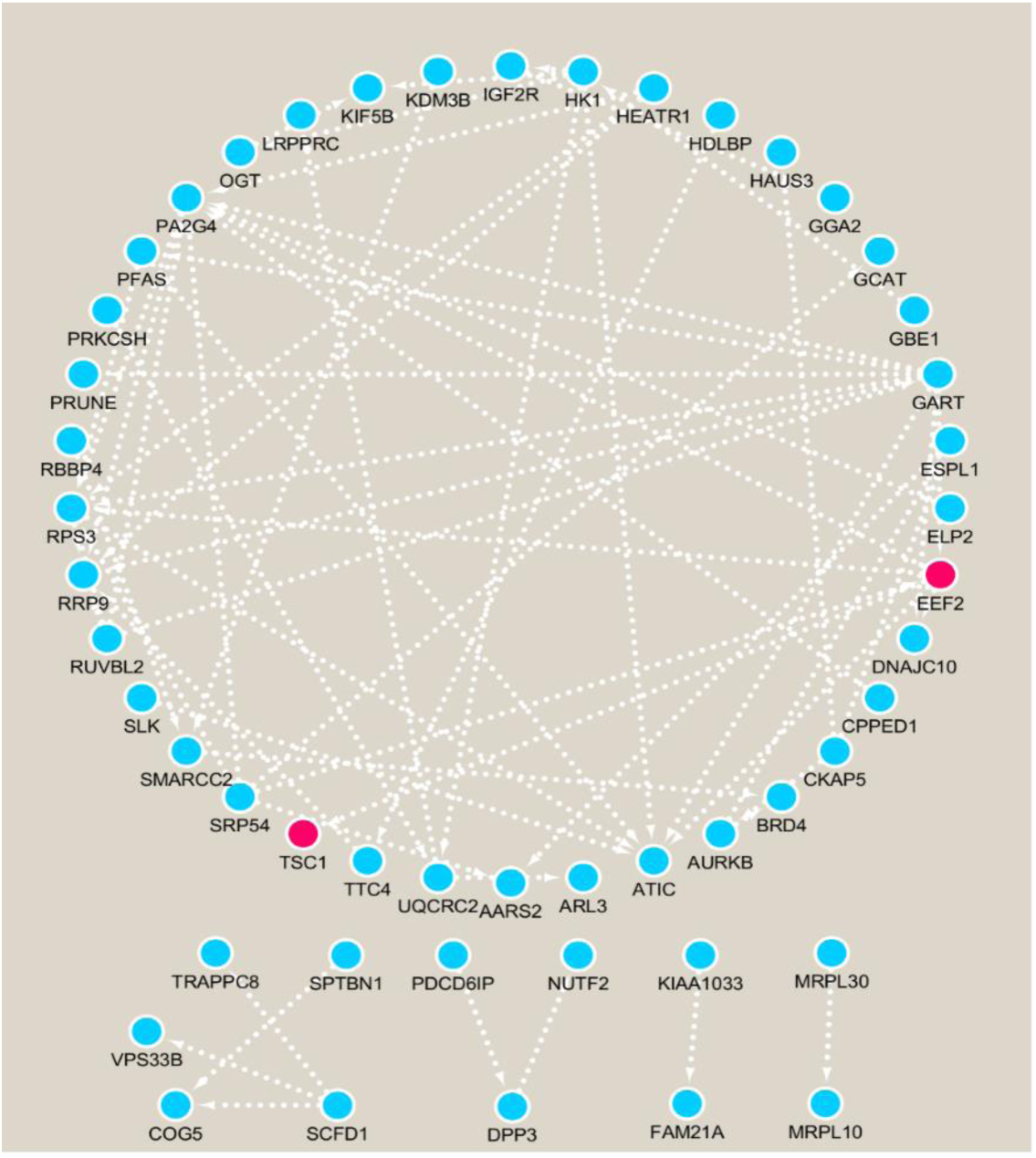
PPI network of 55 mutant proteins, red node represents driver gene, blue node represents passenger gene.

**Figure 5.**
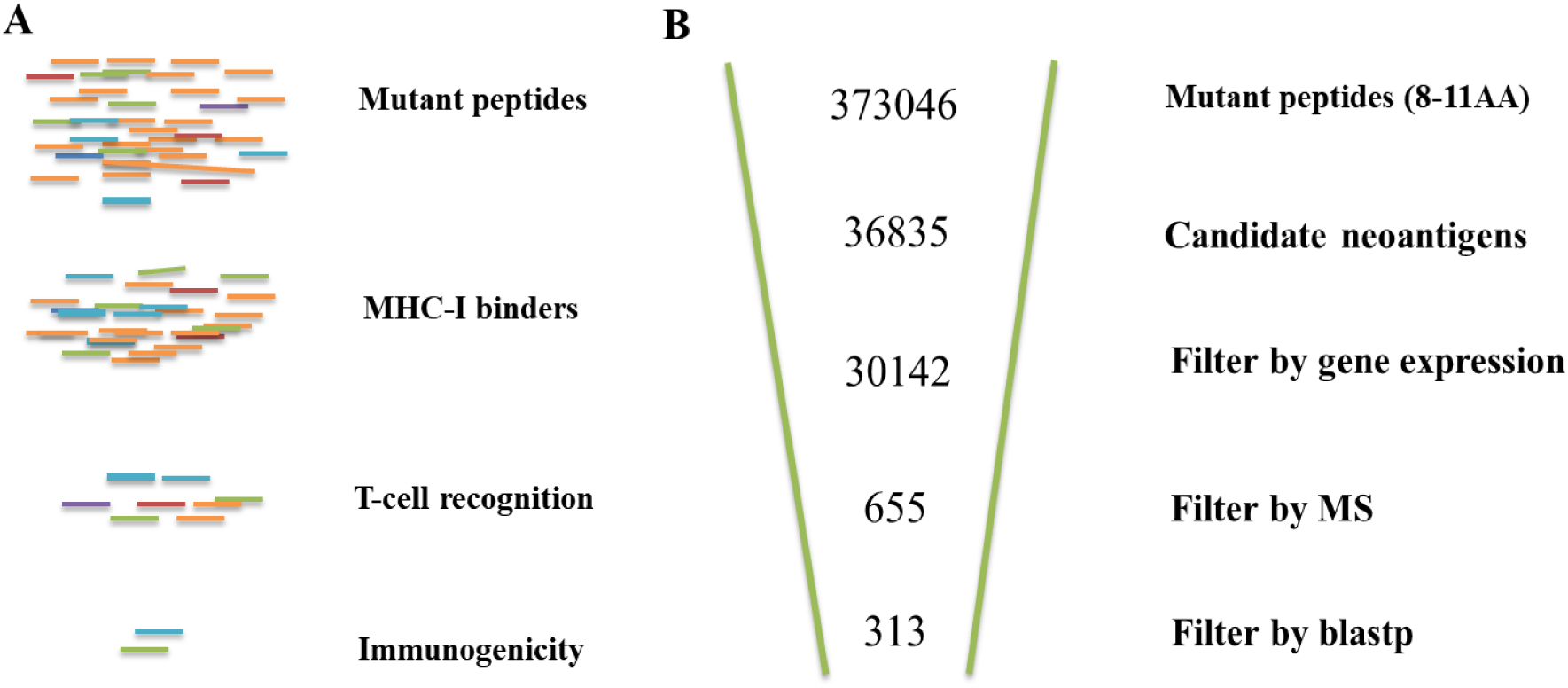
**(A)** Only a small minority of the peptides are capable of eliciting CD8+T cell responses among potential neopeptides. **(B)** Results overview for the neoantigen discovery by proteogenomic workflow: ProGeo-neo.

So far, the large body of publicly available MS/MS data has been used for training HLA-I binding prediction [44, 48, 49] or building spectral libraries [50]. Mass spectrometry analyses of peptides from the peptide-human leukocyte antigen(HLA) complex have enabled the discovery of HLA ligandome tumor antigens for personalized vaccines, LC-MS/MS being an important part for identification of potential cancer neoantigens. Therefore, proteogenomic method was used to predict personalized neoantigens in this study. To the best of our knowledge there is currently no publicly available integrated computational pipeline to perform prediction of neoantigens in consideration of proteomics data. Our ProGeo-neo presents a bioinformatic pipeline for mining tumor specific antigens from next-generation sequencing including genomic and mRNA expression data, incorporating latest netMHCpan (v.4.0) to predict the binding information of mutant peptides to class I MHC molecules, achieving MHC-peptides validation at the peptide/protein level by MaxQuant, and checking potential of T-cell-recognization by adding sequencing screening methods. Our approach efficiently captured peptides generated by missense variants. With massively parallel sequencing and proteomics becoming increasingly applicable, such analyses will be increasingly useful for cancer research and immunotherapy.

Historically our group utilized proteogenomics strategy for genome reannotation [51, 52], we also established methods to study tumor fusion genes and virus genome insertion into human tumor genome by proteogenomics analyses [53, 54]. In recent years the proteogenomics strategy has gained wider attention in high-profile cancer research, because of its integration nature to directly connect proteomics data and genomics data [55, 56]. However, using proteogenomics method to discover neoantigen is still a new field in its infancy. The main bottle neck lies in the lower coverage of proteomics peptide detection compared to whole-genome sequencing, especially when neoantigens are among the low-abundance proteins and hardly detectable by general mass spectrometry. The keep-progressing development of high accuracy tandem mass spectrometry technology has greatly improved the peptide capture coverage and depth, yet the most ideal proteomics data for proteogenomics discovery of neoantigens are HLA-binding enriched peptidomics data. As such data are being deposited into public resources [57], researches in this field are anticipated to increase, and our pipeline ProGeo-neo would be a useful bioinformatics aid.

We are aware that our study bears several limitations. First, mutations in tumor include not only point mutations but insertion/deletion mutations or frame shift mutations. It is clear that insertion/deletion and frameshift mutation will lead to larger changes in amino acid sequence and spatial structure, which may also generate potential neoantigens, these genomic variations are not yet considered in current version of our tool. A recent study reported that noncoding regions are the important source of targetable tumor-specific antigens [58]. Neoantigen discovery currently relies on whole exome sequencing and predominantly on the predictive power of algorithms that infer HLA-ligand binding. Thus, focusing on the exome as the only source of tumor-specific antigens is very restrictive. Lastly, our work does not currently address prediction of HLA class II binding epitopes presented to CD4+T cells because so far the prediction accuracy for MHC-II binders is still low due to their longer size. These represent future developing directions for us, as well as for the field of proteogenomics prediction and selection of personalized tumor neoantigens, which would help the important vaccine-based immunotherapy for human cancers.

## 5. Conclusions

Mass spectrometry-based proteogenomics is a promising new strategy for identifying tumor neoantigens. We propose a workflow to predict and verify HLA-I binding neoantigen peptides on personalized level with proteogenomic methodology. The workflow is constructed based on the genomics and proteomics data of Jurkat leukemia cell line but suitable for predicting neoantigens in other individual solid tumors. The workflow is implemented in a software package, called ProGeo-neo. ProGeo-neo can be applied not only to identify putative neoantigens, but also to screen neoantigens for their immunogenicity. The neopeptides identified from this customized database workflow are of much higher quality as compared to those identified using genomic data only, and could greatly reduce the validation scope of potential subsequent experiments. With parallel sequencing and proteomics becoming increasingly applicable, ProGeo-neo may prove to be a useful bioinformatics tool for cancer research and immunotherapy.

## Acknowledgements

Not applicable.

## Availability of data and materials

All data generated or analyzed during this study are included in this published article and its supplementary data files.

## Ethics approval and consent to participate

Not applicable.

## Consent for publication

Not applicable.

## Author Contributions

YL collected and analyzed the data. YL, GW, XT and QL interpreted the results. YL and JO developed the ProGeo-neo. MZ, XS, QL, LC and LX provided constructive suggestions and discussions during the project. YL and LX wrote the manuscript. LX conceived the idea, planned and coordinated the entire project. All authors read and approved the final manuscript.

